# Analysis of the Confidence in the Prediction of the Protein Folding by Artificial Intelligence

**DOI:** 10.1101/2023.05.17.540933

**Authors:** Paloma Tejera-Nevado, Emilio Serrano, Ana González-Herrero, Rodrigo Bermejo-Moreno, Alejandro Rodríguez-González

**Affiliations:** ETS Ingenieros Informáticos, Universidad Politécnica de Madrid, Madrid, Spain; Centro de Tecnología Biomédica, Universidad Politécnica de Madrid, Pozuelo de Alarcón, Madrid, Spain; Margarita Salas Center for Biological Research (CIB-CSIC), Spanish National Research Council, Madrid, Spain

**Author notes:** {, } { } { }.

**Keywords:** Protein Structure Prediction, Machine Learning Metrics, Model Confidence

## Abstract

The determination of protein structure has been facilitated using deep learning models, which can predict protein folding from protein sequences. In some cases, the predicted structure can be compared to the already-known distribution if there is information from classic methods such as nuclear magnetic resonance (NMR) spectroscopy, X-ray crystallography, or electron microscopy (EM). However, challenges arise when the proteins are not abundant, their structure is heterogeneous, and protein sample preparation is difficult. To determine the level of confidence that supports the prediction, different metrics are provided. These values are important in two ways: they offer information about the strength of the result and can supply an overall picture of the structure when different models are combined. This work provides an overview of the different deep-learning methods used to predict protein folding and the metrics that support their outputs. The confidence of the model is evaluated in detail using two proteins that contain four domains of unknown function.

## 1 Introduction

Protein folding refers to the mechanism through which a polypeptide chain transforms into its biologically active protein in its 3D structure, and it has a significant impact on different applications, e.g., drug design, protein-protein interaction, and understanding the molecular mechanism of some diseases.

There are classic methods to determine the structure of a protein, such as X-ray, NMR, and EM. These methods can be costly and time-consuming because they require significant resources and expertise. Protein folding is a complex process that is challenging for different reasons, including a large number of possible conformations, the crowded cellular environment, and the complex energy landscape required to reach the final structure. The emergence of deep learning methods for predicting protein folding has revolutionized traditional biochemistry. These methods enable in silico predictions, followed by laboratory validation of the findings, offering a new approach to the field.

The Critical Assessment of Protein Structure Prediction (CASP) experiments aim to evaluate the current state of the art in protein structure prediction, track the progress made so far, and identify areas where future efforts may be most productively focused. The bi-annual CASP meeting has shown that deep learning methods like AlphaFold, Rosetta, RoseTTAFold and trRosetta, are more effective than traditional approaches that explicitly model the folding process.

AlphaFold2 was introduced as a new computational approach that can predict protein structures with near-experimental accuracy in most cases. The artificial intelligence system, AlphaFold, was submitted to the CASP 14 competition as AlphaFold2, using a completely different model from the previous AlphaFold system in CASP13 [1]. A multimer version has been released, which allows scientists to predict protein complexes and detect protein-protein interactions. The use of AlphaFold-Multimer leads to improved accuracy in predicted multimeric interfaces compared to the input-adapted single-chain AlphaFold, while maintaining a high level of intra-chain accuracy [2].

To fully utilize these methods, researchers require access to powerful computing resources, which is why alternative platforms have been developed. ColabFold is one such platform that offers an accelerated prediction of protein structures and complexes by combining the fast homology search of MMseqs2 with AlphaFold2 [3]. From early 2022, the Galaxy server is able to run AlphaFold 2.0. as part of the tools offered for bioinformatics analysis. DeepMind and EMBL’s European Bioinformatics Institute (EMBL-EBI) did indeed collaborate to create AlphaFold DB, which offers open access to over 200 million protein structure predictions, with the goal of accelerating scientific research [1, 4]. CAMEO (Continuous Automated Model EvaluatiOn) is a community project, developed by the Computational Structural Biology Group, at the SIB Swiss Institute of Bioinformatics and the Biozentrum of the University of Basel. CAMEO is a service that continuously evaluated the accuracy of protein prediction servers based on known experimental structures released by the PDB. Users can submit several models for a target protein in CAMEO, and it will evaluate up to 5 models. Through CAMEO, Robetta, a protein prediction service, undergoes continual evaluation.

While deep learning models can achieve impressive accuracy on a wide range of tasks, it’s important to carefully evaluate and interpret their predictions to ensure they are reliable and useful. This involves not only selecting appropriate metrics, but also understanding the underlying assumptions and limitations of the model, and how these can impact its predictions. The paper is organized in the following order: Section 2 provides a description of the metrics and scores, while Section 3 describes the tools and resources used in this paper. In Section 4, the results obtained are presented and discussed in Section 5. Finally, Section 6 presents the conclusions of this work and outlines future directions for research.

## 2 Metrics and Scores

It is difficult to understand the reason for the outputs obtained when using deep learning models. There are some common metrics to evaluate the models and choosing the appropriate metric will depend on the application. Interpreting the predictions of a machine learning model is an iterative process, involving ongoing analysis and refinement of the model. By continually evaluating the model’s output, identifying areas of weakness or uncertainty, and refining the model’s parameters and architecture, researchers and practitioners can develop more accurate and reliable models for a wide range of applications.

The GDT (Global Distance Test) is a metric used to measure the similarity between two protein structures with identical amino acid sequences but different tertiary structures. It is primarily used to compare protein structure predictions to experimentally determined structures, and the GDT_TS score, which represents the total score, is a more accurate measurement than the commonly used root-mean-square deviation (RMSD) metric. To calculate the GDT score, the model structure is evaluated for the maximum set of alpha carbon atoms of amino acid residues that are within a specific distance cutoff from their corresponding positions in the experimental structure. The GDT score is a crucial assessment criterion in the Critical Assessment of Structure Prediction (CASP), a large-scale experiment that evaluates current modelling techniques and identifies primary deficiencies [5].

The LDDT (Local Distance Difference Test) is a score that measures the local distance differences between all atoms in a model, regardless of their superposition, to evaluate the plausibility of stereochemistry [6]. AlphaFold2 reports the predicted quality of a protein as a per-residue pLDDT score, which is used to assess intra-domain confidence [1]. The pLDDT score ranges from 0 to 100, with higher scores indicating higher quality predictions. Generally, residues with pLDDT scores greater than 90 are considered reliably predicted, while scores between 70-90 are less reliable, and scores below 50 are low quality. AlphaFold2 calculates the pLDDT score by comparing the predicted distances between pairs of atoms in the predicted protein structure with the corresponding distances in a reference set of experimentally determined protein structures. This comparison is done at the level of individual residues, and the resulting score reflects the similarity between the predicted and reference structures at each residue position.

The Predicted Aligned Error (PAE) is a metric used by the AlphaFold system to assess the quality of predicted protein structures. PAE measures the average distance between the predicted and true residue positions in a protein structure, after aligning the predicted structure with the true structure. First, the predicted protein structure is aligned to the true structure using a variant of the Kabsch algorithm, which finds the optimal rotation and translation that aligns the predicted structure to the true structure. Next, the distance between each predicted residue position and its corresponding true residue position is calculated. Finally, the average distance between the predicted and true residue positions is computed, resulting in the PAE score. The Predicted Aligned Error (PAE) is best used for determining between domain or between chain confidence. Like pLDDT, PAE is reported as a per-residue score, with higher scores indicating better prediction accuracy. Residues with PAE scores below a certain threshold are outliers and are often indicative of regions of the protein that were difficult to predict or where the predicted structure deviates significantly from the true structure. The template modelling score (TM-score) is developed for automated evaluation of protein structure template quality [7]. The predicted TM-score (pTM-score) takes into account the probability of each residue being resolved by weighting its contribution accordingly.

The present study examined two proteins containing four domains of unknown function (DUF1935). ARM58 is an antimony resistance marker found in *Leishmania* species [8, 9], while ARM56 has orthologues in *Trypanosoma* spp. but does not confer antimony resistance [9, 10]. The protein sequences were inputted into various prediction methods, including tr-Rosetta, tr-RosettaX-Single, RoseTTAFold (using Robetta), ColabFold, AlphaFold2 (Galaxy server), and the AlphaFold Protein Structure Database. By comparing and optimizing the results from these different methods, the study aimed to determine the most effective methodology for utilizing deep learning techniques in predicting protein structure. Such techniques can serve as both routine and complementary tools, providing guidance and accelerating experimentation. The diverse outputs obtained from different protein prediction models suggest that discrepancies between the models could reveal valuable information about the flexibility of the proteins. Careful observation of these variations is necessary to determine their significance and relevance for understanding protein structure and function.

## 3 Material and Methods

ARM58 and ARM56 are encoded by the genes LINFJ_34_0220 and LINFJ_34_0210, respectively. The protein sequences contain four domains of unknown function (DUF1935), which were downloaded in FASTA format from UniProt. Predicted AlphaFold protein structures for ARM58 and ARM56 were also downloaded from the Protein Structure Database, including the PDB file and the predicted aligned error. The Galaxy server was used to generate AlphaFold 2.0. predictions for both proteins. It was also used ColabFold v1.5.1. [3] which utilizes MMseqs2 and HHsearch for sequence alignments and templates. The protein structure prediction service Robetta was used to obtain structures using RoseTTAFold [11]. Additionally, the trRosetta server [12, 13] was used for protein structure prediction by transform-restrained Rosetta. Also, the trRosettaX-Single does not use homologous sequences and templates [14]. Finally, the relax_amber notebook was used to relax the structures using amber, with the outputs labelled as unrelaxed. Protein predictions were visualized using UCSF ChimeraX v1.5 [15].

## 4 Results

ARM58 and ARM56 protein predictions using AlphaFold2 were downloaded from the Protein Structure DB. ARM58 and ARM56 contain 517 and 491 amino acids, respectively. AlphaFold2 was also run on the Galaxy server. Both prediction outputs were compared using ChimeraX, by matching both structures (figure 1 A, B). The PAE plot indicates the expected distance error for each residue position where the low-error squares correspond to the domains for both proteins (figure 1 C, D).

**Figure 1.**
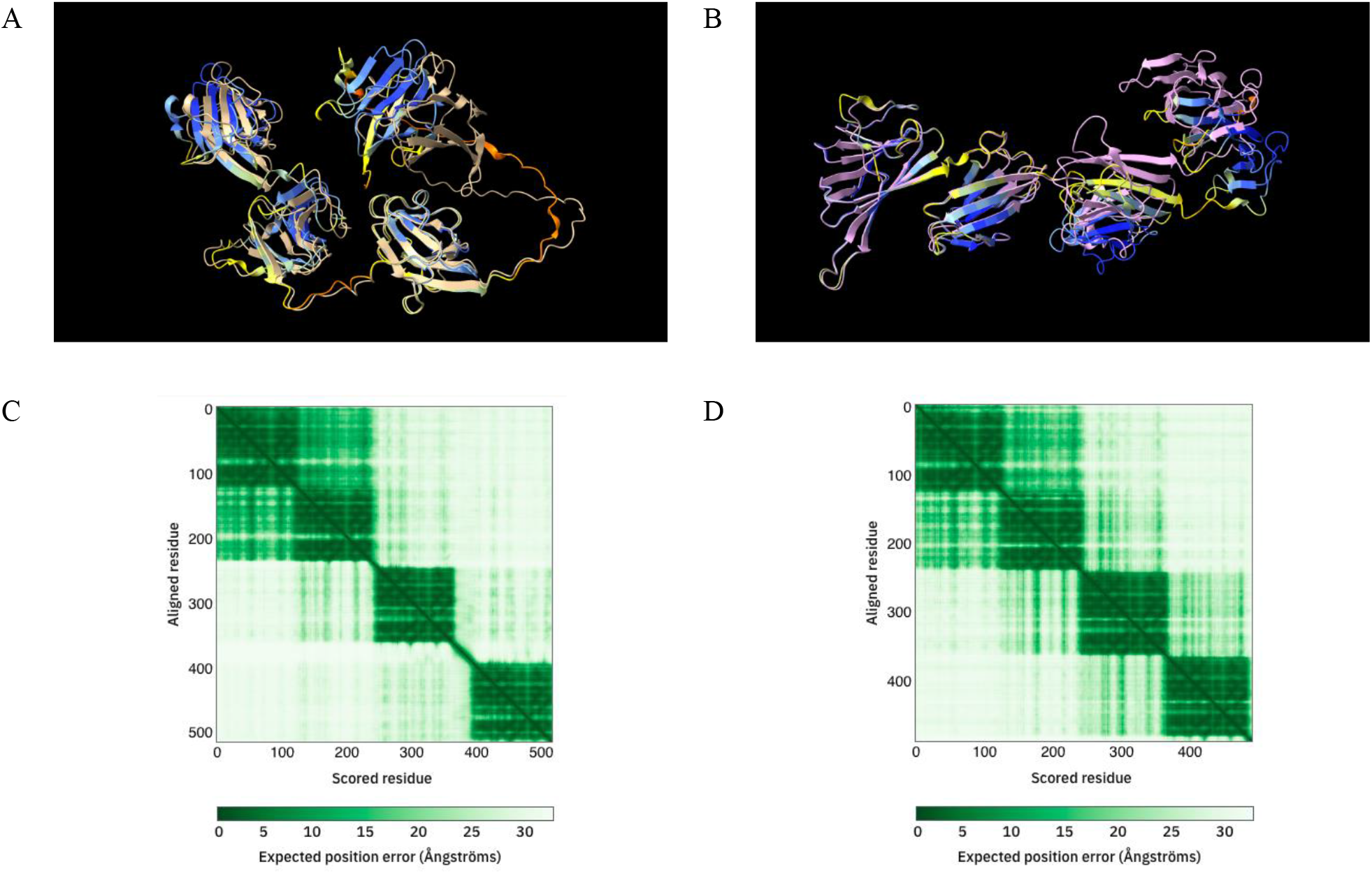
Protein structure prediction for ARM58 and ARM56. Predicted per-residue for the best model from AlphaFold DB coloured with the predicted aligned error (dark blue (100 - 90): high accuracy expected; light blue - yellow (90 - 70): expected to be modelled well; yellow - orange (70 - 50): low confidence; orange – red (50 - 0): may be disordered) and AlphaFold2 prediction best model obtained in Galaxy (brown and purple) for ARM58 (A) and ARM56 (B). Protein prediction structures are visualized using ChimeraX. Predicted aligned error representing the four domains for ARM58 (C) and ARM56 (D) from AlphaFold protein structure database.

Next, ColabFold using MMseqs2 was used to perform an analysis on the metrics and outputs. The tool was run three times with the default parameters generating small variations for pLDDT and pTM (table 1).

**Table 1.** Average and Standard deviation from the metrics pLDDT and pTM obtained in ColabFold using MMseqs2 for ARM58 and ARM56, running the protein structure predictions three times.

|  |  | ARM58 | ARM56 |
| --- | --- | --- | --- |
| pLDDT | Average | 80.433 | 86.733 |
|  | Standard deviation | 0.577 | 4.619 |
| pTM | Average | 0.410 | 0.486 |
|  | Standard deviation | 0.003 | 0.004 |

Out of the three different runs, two protein prediction structures were selected based on their differing pLDDT and pTM scores for additional examination. The selected structures were visualized using ChimeraX software and color-coded based on the pLDDT values (figure 2).

**Figure 2.**
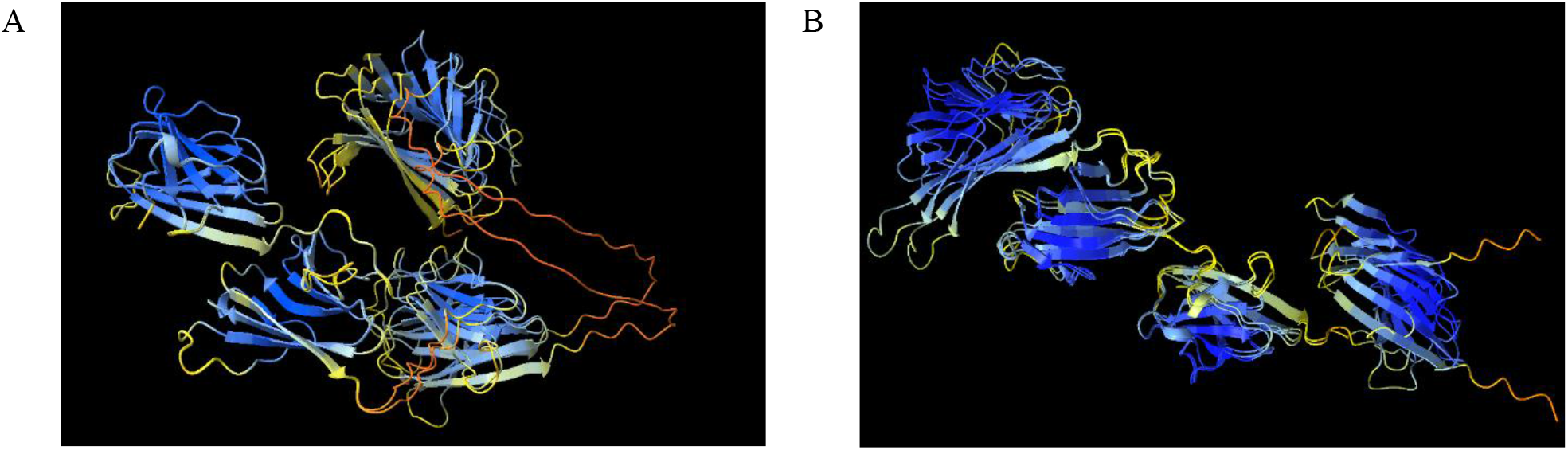
Protein structure prediction for ARM58 (A) and ARM56 (B) using ColabFold (MMseqs2). Two resulting structures were coloured with the model confidence (dark blue: pLDDT > 90; light blue: 90 > pLDDT > 70; yellow: 70 > pLDDT > 50; orange: pLDDT < 50). Protein prediction structures were visualized using ChimeraX.

The trRosetta server, along with trRosettaX-Single, was used to predict the protein structures of ARM58 and ARM56. The models obtained using trRosetta have high confidence, with estimated TM-scores of 0.608 for ARM58 and 0.650 for ARM56. Moreover, the confidence of the models generated using trRosettaX-Single was low, with estimated TM-scores of 0.275 for ARM56 and 0.266 for ARM58. The predicted per-residue LDDT values for the best model reflected the lower confidence per residue in the tr-RosettaX-Single models (figure 3).

**Figure 3.**
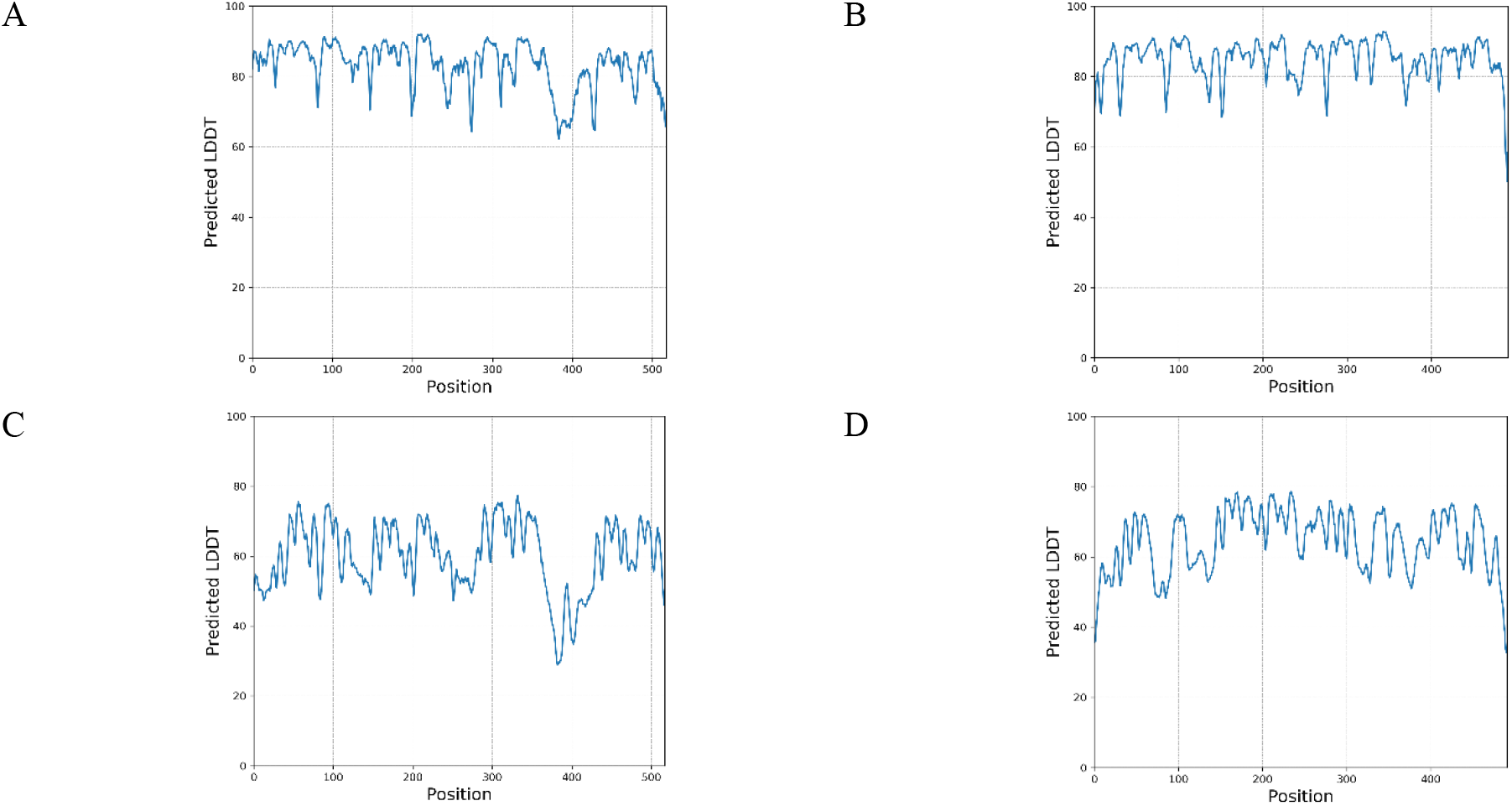
Predicted per-residue for the best model in ARM58 and ARM56. Predicted LDDT from tr-Rosetta (A) and trRosettaX-Single (C) for ARM58. Predicted tr-Rosetta (B) and tr-RosettaX-Single (D) for ARM56.

Eventually, the structures of ARM56 and ARM58 were predicted using the modelling method RoseTTAFold via Robetta service. The predicted Global Distance Test (GDT) confidence scores for the models were 0.79 for ARM56 and 0.76 for ARM58. Finally, the tr-Rosetta and RoseTTAFold prediction models were slightly different from those previously described. In the end, the tr-Rosetta model was compared to the most similar of the five models predicted with RoseTTAFold (figure 4).

**Figure 4.**
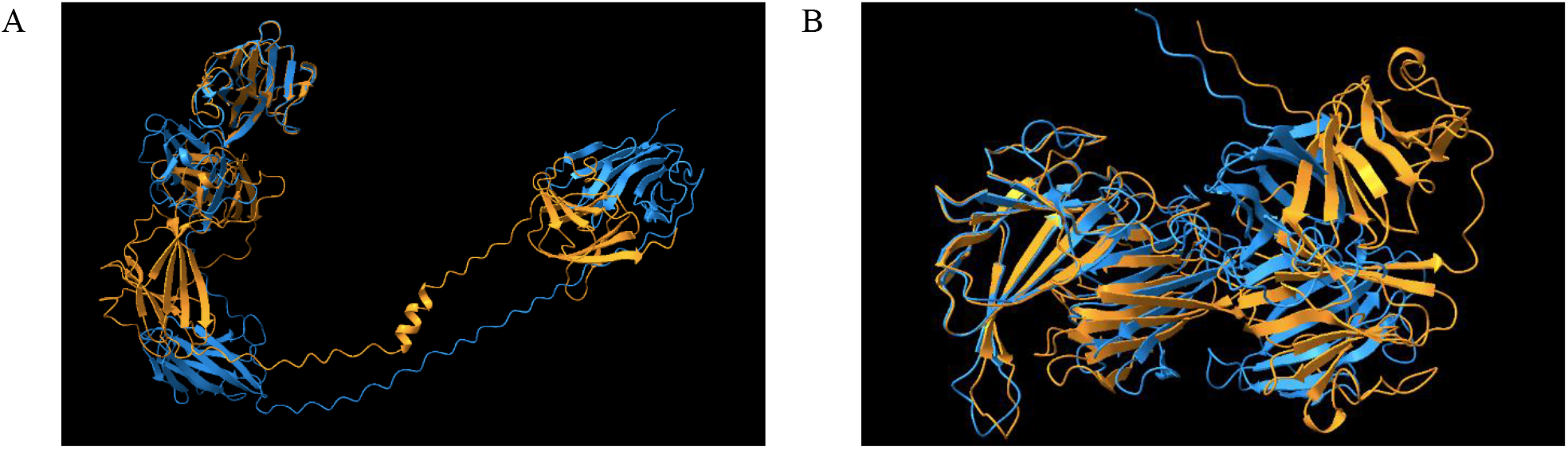
Predicted per-residue for the best model in ARM58 (A) and ARM56 (B) using tr-Rosetta (blue) and RoseTTAFold (orange). Protein prediction structures were visualized using ChimeraX.

## 5 Discussion

The unknown exact positions of atoms in a protein can be related to flexible regions because the flexibility or rigidity of a protein is determined by the relative positions of its constituent atoms. The flexibility of certain regions in a protein can be essential for its function, as they allow for conformational changes that are needed for the protein to interact with other molecules or carry out its biological role [16]. In the present work, ARM58 and ARM56 predicted models were able to define the beta sheets present in each domain (figure 1). However, the exact positions of the atoms between the four domains are not fully defined (figure 2). This could be related to flexibility, or it could be that the exact positions are uncertain. When regions are extensive and have a pLDDT value below 50, they exhibit a strip-like visual pattern and ought to be consider a projection of disorder, rather than an indication of actual structure [17].

Protein prediction models may generate structures that deviate from those produced by traditional models. Explaining these models can be challenging, therefore, it is important to interpret the metrics correctly. The accuracy of the models is given by the pLDDT, PAE, or pTM scores, which need to be analysed by researchers. There are fast and reliable prediction methods such as ColabFold [3] that provide accurate protein prediction structures. However, variations in the metrics (table 1) can lead to diverse outcomes. Some models can work better on a specific set of proteins. In this case, trRosetta and trRosettaX-Single show notable differences in the confidence given by the pLDDT score (figure 3). Each method generates different metrics for evaluating confidence; for instance, RoseTTAFold, when used with Robetta, provides the predicted GDT (Global Distance Test) confidences. Distinct structure prediction systems generate varying models (figure 4), making modelling a challenging task.

A combination of all the structures, evaluated through various metrics, can provide complementary information about the structures of proteins, especially in cases where no prior information is available from X-ray, NMR or EM techniques. The proteins ARM58 and ARM56 each consist of four domains whose functions are unidentified. It is unknown the mechanism that leads to antimony resistance to *Leishmania* spp upon ARM58 overexpression. Here, different protein prediction methods have been used to compare the predicted structure of ARM58 and ARM56. There are differences between models, and each model can generate variations in each run. Additionally, alternative installations could also produce different outputs. By accurately simulating a variety of temporary protein complexes, end-to-end deep learning underscores opportunities for future enhancements that can enable dependable modelling of any protein-protein interaction that researchers want to explore [18]. Factors such as amino acid sequence composition or protein function could provide more information. In addition to predicting protein structures, AlphaFold2 also provides insights into the flexibility of residues, or protein dynamics, that are encoded in these structures [19].

## 6 Conclusions and Future Work

Proteins can indeed be represented as graphs or networks, depending on the specific context. For example, in the case of protein-protein interaction (PPI), each protein can be represented as a node in a network, and the links between nodes would indicate the interactions between the corresponding proteins. In the case of protein structure representation, each amino acid in the protein can be represented as a node, and the edges between nodes would represent the distances between them. Protein structure prediction also enables the determination of protein-ligand interactions. However, protein regions that lack specific structures make it difficult to determine the exact positions of atoms. Consequently, variations in these positions could result in different possible structures, reflecting the protein’s flexibility. Protein folding is a critical process in the development of new drugs, as the three-dimensional structure of a protein determines its function and interactions with other molecules. Thus, it is important to determine which prediction provides more information about the structure in order to obtain insights into different functional states or biologically relevant features.

There is a need to understand the generation of different outputs through deep learning models. When there is complementary material from classic prediction methods or experimental assays, it is possible to obtain additional information. Frequently, there is no additional investigation previously done, and therefore, it is necessary to rely on the metrics and understand the predictions, especially when further experiments will be performed.

## Acknowledgments

This work is a result of the project “Data-driven drug repositioning applying graph neural networks (3DR-GNN)”, that is being developed under grant “PID2021-122659OB-I00” from the Spanish Ministerio de Ciencia e Innovación. This work was funded partially by Knowledge Spaces project (Grant PID2020-118274RB-I00 funded by MCIN/AEI/10.13039/501100011033).

